# Gene expression signatures underlying inbreeding depression as revealed by whole-transcriptome analysis of selectively bred strains of the Pacific Oyster (*Crassostrea gigas*)

**DOI:** 10.1101/2022.05.01.490184

**Authors:** Chenyu Shi, Fuqiang Zhang, Qi Li, Shikai Liu

## Abstract

Exploring molecular mechanisms of inbreeding depression is significant for the conservation and sustainable use of the rare breed with a specific trait of high production value. In this work, we conducted whole-transcriptome analysis of two selectively bred Pacific oyster strains with one inbred strain showing significant growth depression. A total of 7980, 2677, and 28 differentially expressed protein-coding genes (DEGs), lncRNAs (DELs), and miRNAs (DEMs) were identified, respectively. The majority of DEGs and predicted target genes of DELs and DEMs were significantly enriched in biological process including immune response, cell proliferation, and apoptosis pathways. A set of genes with specific expression patterns as revealed by cluster profiling and enrichment analysis were identified, which may underlie inbreeding depression of the inbred strain. Furthermore, a competing endogenous RNA (ceRNA) network was constructed by integrative analysis of DEGs, DELs, and DEMs, supporting that ncRNAs, as regulators of gene expression, could be critical modulators in multiple subsystems involved in inbreeding depression.

## Introduction

Selective breeding toward genetic improvement of production and performance traits is an efficient way to meet increasing demand of food. In the post-genomics era, the further understanding of biotic mechanisms driving the species-traits formation and rapid development of breeding-related technologies simplify the breeding process and provide new solutions and opportunities for rapid and effective selective breeding. In this case, the rare breeds with a specific trait of high production value have an immense widespread interest and the conservation or development of rare breeds is gradually becoming an active research field (Molnár et al. 2014; Bowley et al. 2019; Nandolo et al. 2019). However, a breeding program with rare breeds as the founder population develops opportunities and faces challenges. A loss of genetic variability and an increase in inbreeding depression because of rare breeds’ small effective population size (Ne) and an effect of genetic drift on breed characteristics (Danchin-Burge et al. 2010). Thus, exploring and appreciating the biochemical effects and causes of inbreeding depression is significant for the conservation and sustainable use of the rare breed.

A lot of research shows that inbreeding depression is associated with a large number of gene transcripts, primarily genes related to metabolism, stress, and defense (Paige 2010). Apart from this, accumulating evidence indicates that the epigenetic mechanisms that can affect phenotype from one generation to the next are also important, such as DNA methylation, histone modifications, and non-coding RNAs (ncRNAs) (Triantaphyllopoulos et al. 2016). ncRNAs are transcripts that do not produce any functional peptides or proteins, but play an essential role in regulating gene expression at multiple levels. Long non-coding RNAs (lncRNAs) and microRNAs (miRNAs) are the two prominent families of ncRNAs that have gained significant attention as regulators of gene expression and epigenetics (Kaikkonen et al. 2011). Despite their great function in regulating various biological processes, non-coding transcripts are much less understood than protein-coding transcripts, especially in aquaculture species. Systematic identification of ncRNAs is limited to a few species. Most of the relationships between ncRNAs and their potential target genes are still unclear in this new area of research (The Aquaculture Genomics, Genetics and Breeding Workshop et al. 2017).

Molluscan shellfish is a major product of world aquaculture, and many selective breeding programs for oyster, clam, scallop, and mussel species have been initiated worldwide (Fang et al. 2021). In our previous breeding practice of the Pacific oyster, a rare orange-shell variant line of *C. gigas* was obtained based on four variant orange-shell oysters and through several generations of family selection for orange shell color and growth trait (Han et al. 2019). The orange-shell oyster line is a typical inbred line, and related assessment shows that though the genetic variability of this orange line was maintained over generations of intense mass selection, inbreeding depression may strongly affect its viability and growth trait (Fang, Han, and Li 2021; Han and Li 2018). Therefore, the orange shell line is an ideal inbred animal model to study the molecular mechanism of inbreeding depression in rare breeds.

To comprehensively identify and analyze the whole transcriptome expression profile in the orange-shell variant line of *C. gigas*, we selected the black-shell oyster line for comparison, which is the base source of the rare orange-shell variant line. Our study was the first to identify and characterize the lncRNAs and miRNA in *C. gigas* from whole tissue at the genomic level. Differential protein-coding transcripts expression was characterized to gain additional insight into the potential difference of two oyster lines’ genetic regulation. Furthermore, we determined expression profiles and sequence characters of ncRNAs and predicted their potential target genes to analyze their functions. We also systematically constructed a lncRNA-associated ceRNA network by combining bioinformatics and correlation analyses to find the potential hub lncRNAs. These results will provide helpful information for further investigation of molecular mechanisms of inbreeding depression in rare breeds and promoting molecular breeding in the Pacific oyster.

## Materials and Methods

### Experimental Oyster and Sample Collection

In this study, the basic black-shell (HH) and the orange-shell variant (CC) *C. gigas* lines of full-sib families used were established on the same day. Three months later, the Pacific oysters from two lines were collected randomly. Nine individuals from the black-shell family were larger (“HH” group, mean shell height, 18.60 ± 0.21 mm), and nine from the orange-shell family were smaller (“CC” group, mean shell height, 14.20 ± 0.19 mm). In each group, all nine oysters of the soft part were dissected an d mixed together in threes, creating three biological replicates for each group. Isolated tissues were flash-frozen in liquid nitrogen. Offshore farms for Pacific oyster sample collection were maintained by the College of Fisheries, Ocean University of China (OUC), Qingdao, China. All animal care and use procedures were approved by the Institutional Animal Care and Use Committee of Ocean University of China (Permit Number: 20141201) and were performed according to the Chinese Guidelines for the Care and Use of Laboratory Animals (GB/T 35892-2018).

### RNA extraction, library construction and sequencing

Total RNA from two lines were extracted respectively using TRIzol reagent (Invitrogen, Carlsbad, CA, USA) according to the manufacturer’s protocol. The purity, concentration, and integrity of the total RNA were measured by NanoDrop 2000 (Thermo Fisher Scientific) and RNA Nano 6000 Assay Kit of the Bioanalyzer 2100 system (Agilent Technologies, CA, USA). Approximately 3 μg of total RNA per sample was used for RNA library construction. Following the manufacturer’s recommendations, Strand-specific RNA sequencing library and small RNA sequencing libraries were generated respectively using NEBNext^®^ Ultra™ RNA Library Prep Kit and NEBNext^®^ Multiplex Small RNA Library Prep Set for Illumina^®^ (NEB, Ipswich, MA, USA). After the cDNA synthesis and PCR amplification, the sequencing of each strand-specific Libraries and small RNA libraries was performed on Illumina HiSeq 2500 platform and raw reads were generated.

### Bioinformatics pipeline for isolation and identification of lncRNAs and microRNAs

#### Identification of lncRNAs

The pipeline used for the identification of lncRNA has been described in Figure S1-A. After examining the quality of raw data using FastQC (v0.11.8), quality control and reads statistics of Raw data were determined by fastp (v0.22.0) (Chen et al. 2018). The remaining clean reads were subsequently aligned to the *C. gigas* reference genome (https://www.ncbi.nlm.nih.gov/genome/10758) using HISAT2 (v2.1.0) (Kim et al. 2019). The mapped reads of each sample were assembled by using StringTie (v2.1.4) (Pertea et al. 2015). Then, the assembled transcripts were annotated using the gffcompare (v0.11.2) (Pertea et al. 2020). And the unknown transcripts were remained used to screen for putative lncRNA:

(1) The newly assembled transcripts were selected for the identification of lncRNAs, including that share at least one splice junction with a reference transcript (“j”), those having multi-exon with at least one overlaps (“o”), those exonic overlap on the opposite strand (“x”), those fully contained within a reference intron (“i”) and those located in intergenic regions (“u”);

(2) The transcripts shorter than 200 bp length and less than two exons have been discarded;

(3) The transcripts show no or weak protein-coding potential based on the prediction results of CPC2 (Kang et al. 2017), PLEK (Li et al. 2014), LGC (Wang et al. 2019) and CPAT (Wang et al. 2013);

(4) the transcripts exhibit no significant hit (E-value = 1e-3) according to HMMER3 (Mistry et al. 2013) search against the Pfam protein domain database (release Pfam33.1) (https://pfam.xfam.org/);

(5) transcripts with low abundance (FPKM < 0.5) are discarded.

Finally, the remaining transcripts were considered reliable lncRNAs. The different types of lncRNAs were classified based on their location on the genome, including long intergenic noncoding RNAs (lincRNAs, class code “u”), antisense lncRNAs (class code “x”), intronic lncRNAs (class code “i”), and sense lncRNAs (class code “j, o”).

#### Identification of microRNAs

The pipeline used for the identification of microRNAs has been described in Figure S1-B. The raw data from small RNA sequencing were subject to the FastQC to assess the quality of small RNA sequencing, and then the raw data were filtered by using cutadapt (v3.5) (Martin 2011) and trimmomatic (v0.39) (Bolger et al. 2014) to remove 3’ adapters, and low-quality reads. The clean sequences between 18 and 30 nucleotides in length were obtained. The rest of the sequences were aligned to the *C. gigas* reference genome using bowtie (v1.3.0) (Langmead et al. 2009). Subsequently, the matched clean reads were aligned with Rfam database (https://rfam.xfam.org/) to remove other known non-coding RNA. The filtered reads were aligned with the *C. gigas* repeat and exons sequences to remove mapped parts.

Finally, the remained un-annotated reads were used to predict and identify novel miRNA precursor sequences and structures by mirDeep2 (v2.0.1.3) (Friedländer et al. 2012) pipeline. Additionally, in this analysis, invertebrate miRNAs in miRbase (http://www.mirbase.org/), including *Haliotis rufescens, Lottia gigantea, Melibe leonina, Caenorhabditis elegans*, and *Drosophila melanogaster* were used to improve the prediction performance of miRDeep2. RNAfold was used to confirm the structures of the predicted miRNAs and keep only novel miRNAs with a rand fold P-value ≤ 0.05 and a miRDeep2 score ≥0.

### Differential Expression Analysis and Target gene prediction of DE-lncRNAs and DE-microRNAs

StringTie was used to calculate the expression levels of lncRNAs and mRNAs in Strand-specific Libraries. And the expression levels of miRNAs were calculated by the quantifier.pl module from miRDeep2. Identification of differentially expressed transcripts between two lines was performed using DESeq2 (v1.31.0) (Michael et al. 2014) R package. Adjusted p-value < 0.05 and absolute value of log2(fold change) > 1 were considered as significantly differential expressions of mRNAs (DEGs), lncRNAs (DELs), and miRNAs (DEMs).

Potential target genes of the lncRNAs were predicted based on their regulatory patterns, which were classified into cis-and trans-acting groups. These protein-coding genes transcribed within a 10 kb upstream or downstream of lncRNAs were searched as potential cis target genes by using BEDTools (v2.29.2) (Quinlan and Hall 2010). Furthermore, the trans-acting target genes were determined by calculating the expression correlation between the expression level of lncRNAs and mRNAs. And the correlation in expression was evaluated using Pearson’s correlation coefficient (PCC) (|r| > 0.99 and p < 0.01).

miRNA usually regulates gene expression through binding to the 3’ untranslated region (3’UTR) of target mRNAs. The pairs of miRNAs-lncRNAs or miRNAs-mRNAs were predicted using miRanda (v3.3a) (Enright et al. 2003), RNAhybrid (v2.1.2) (Kruger and Rehmsmeier 2006), and Targetscan software (Lewis, Burge, and Bartel 2005). The overlap of the predicted results from the three programs was considered to represent the final result of predicted target mRNAs. For each prediction method, high efficacy targets were selected by the following criteria: (1) miRanda: total score ≥ 125, total energy ≤ −20 kcal/mol; (2) RNAhybrid: mfe ≤ −20 kcal/mol. Then, the predicted miRNAs-lncRNAs pairs were integrated with the miRNAs-mRNAs pairs by the shared miRNAs. Finally, the interaction network was visually displayed using Cytoscape v3.8.0.

### Genes functional annotation and enrichment analysis

To obtain current functional annotations of the *C. gigas* protein-coding genes, we blast protein-coding genes sequences to three different databases: NCBI non-redundant (nr) protein database, EggNOG, and BlastKOALA by using DIAMOND (v2.0.5.143) (Buchfink, Xie, and Huson 2015), with an E-value cutoff of 1e-3. To have more complete results, we integrated the results for the three annotation strategies and built the custom OrgDb for *C. gigas* by using AnnotationForge (v1.35.1) (Marc Carlson 2017) R package.

Gene ontology (GO) and Kyoto Encyclopedia of Genes and Genome (KEGG) pathway enrichment analysis of DEGs and Target genes of DELs and DEMs by using ClusterProfiler (v4.1.4) (Wu et al. 2021) R package, with a cutoff of adjusted P-adj < 0.05. To reduce the redundancy between enriched GO terms and their associated inference related to DEGs, redundant GO terms with > 95% overlap with similar terms were further clustered using GOMCL (Wang et al. 2020).

## Results

### Identification and Characterization of lncRNAs and miRNAs

Strand-specific RNA-sequencing of the six libraries yielded a total of 380,493 raw paired-end reads with a length of 2×150 bp. And after discarding low -quality, adaptor, and poly-N sequences, about 99.6% (379,138,356) of raw reads remained as clean reads for the subsequent analyses. Moreover, in the six libraries, 78.36%-79.55% of clean reads were mapped to the *C. gigas* reference genome. The summary data output for each sample is listed in Table S1-1.

We used a stringent filtering pipeline to identify lncRNAs. Briefly, each sample’s mapped reads were assembled using StringTie, and we generated a new GTF file containing 122,546 assembled transcripts and 49,347 gene loci. Then, a series of filtering strategies were employed to rule out transcripts (details in Materials and Methods). Finally, a total of 6,350 lncRNA transcripts in 4,980 lncRNA loci were identified. Based on the *C. gigas* genome annotation file, we compared the characteristics of lncRNA and mRNA, including Transcription length, exon number, isoform number, and expression level (Fig.1A-D). We found that compared with known mRNAs, the average transcript’s length of lncRNAs is shorter. lncRNAs contain fewer isoforms and exons than mRNAs. And in this analysis, the average expression level of mRNAs was higher than those of lncRNAs.

**Fig. 1 A-D.**
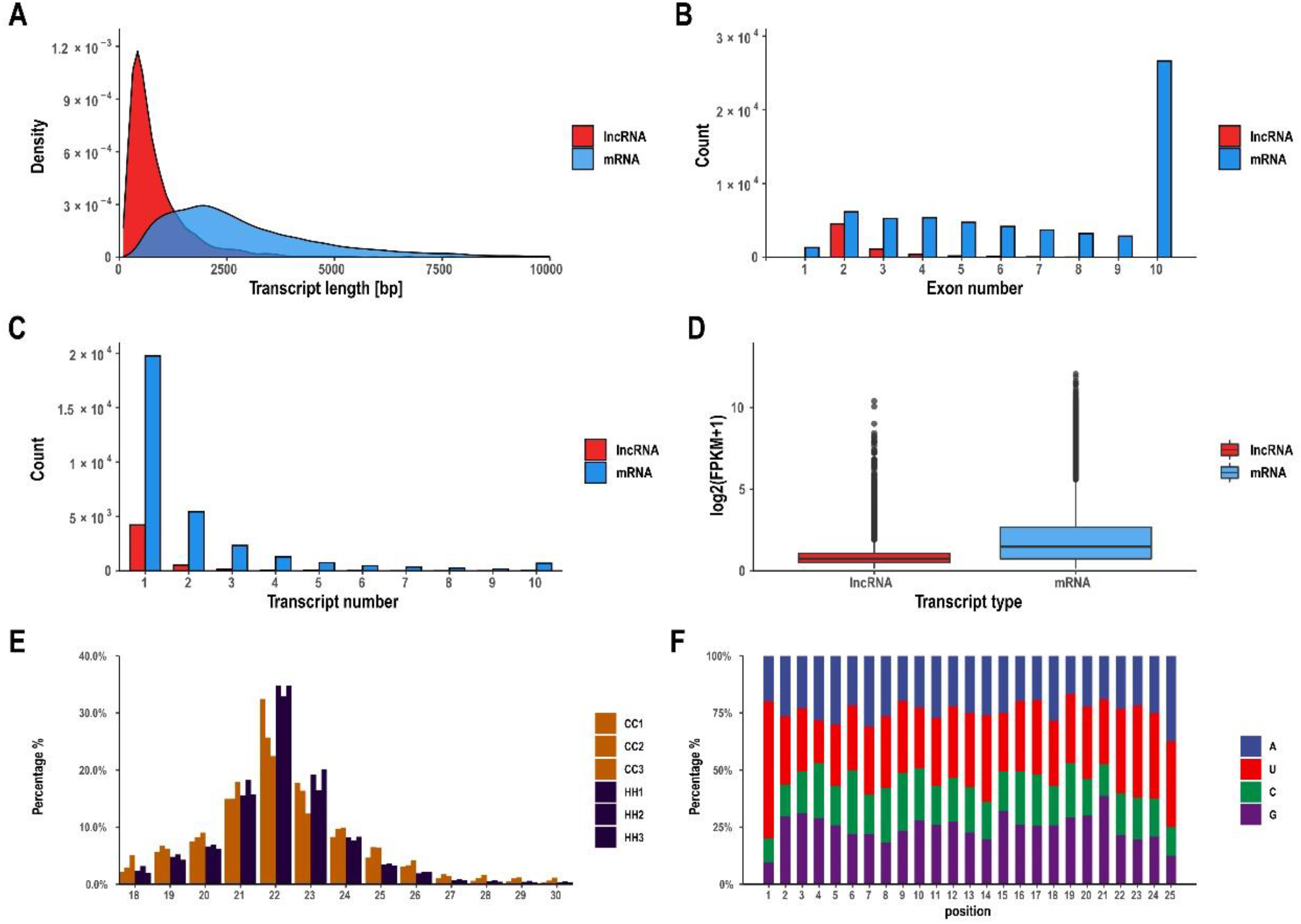
the characteristics of comparing lncRNA and mRNA, including Transcription length(A), exon number(B), isoform number(C) and expression level(D). **Fig.1E** Distribution of small RNAs with different sequence length. **Fig.1F** Nucleotide composition of the predicted miRNAs and their bias positions.

To identify the miRNAs, six small RNA libraries were constructed from the two lines of *C. gigas*. High-throughput sequencing generated a total of 77,087,961 raw reads by high-throughput sequencing (approximately 1.3 million raw reads per library). After removing 3’ adapters and low-quality reads, the clean sequences between 18 and 30 nucleotides in length were obtained for further analysis. The length distribution (Fig 1E) showed that the vast majority of mapped reads are 21-23 nt and reads of 22 nt made up the most abundant group, which is about 63.8% of each library. In order to classify and annotate the small RNAs, all clean reads were mapped against the *C. gigas* reference genome, Rfam database, and the *C. gigas* repeat and exons sequences using Bowtie software (Table S1-2). After filtering out, we identified 376 novel miRNAs by the mirDeep2 pipeline. And by comparing the miRNA precursors sequence with the miRBase database, we found 67 detected miRNAs have high conservation with other species’ miRNAs (Table S2). Figure 1F showed that the first nucleic acid of the predicted miRNAs was enriched with uracil (U). Moreover, we compared with the previous study (Xu et al. 2014) on identifying microRNAs in the Pacific Oyster. We found a high total mapping rate (80.56%) of no mismatch when aligning its mature sequences to hairpin precursor sequences which were predicted in this study.

### Analysis of differentially expressed protein-coding transcripts

Overall, significantly differentially expressed protein-coding transcripts (DEGs), including 4431 up-regulated (55.6%) and 3539 down-regulated (44.4%), were discovered in the HH of CC (Fig S2A and Table S3-1). Then we carried out gene ontology (GO) and pathway enrichment analysis to identify the biological functions of DEGs (Table S4). GO enrichment analyses revealed that DEGs were enriched in 253, 29, and 35 GO terms in biological process, cellular component, and molecular function, respectively. Within the three process categories, complement activation, perisynaptic extracellular matrix, and RNA-DNA hybrid ribonuclease activity were the most represented, respectively (Fig.2A). KEGG pathway analyses showed that DEGs were significantly enriched in 22 KEGG pathways, in which pathways related to apoptosis, axon regeneration, and some conserved immune-related pathways, such as NOD-like receptor signaling pathway, TNF signaling pathway, Toll and Imd signaling pathway, and IL-17 signaling pathway were significantly enriched (Fig.2B).

**Fig.2 A.**
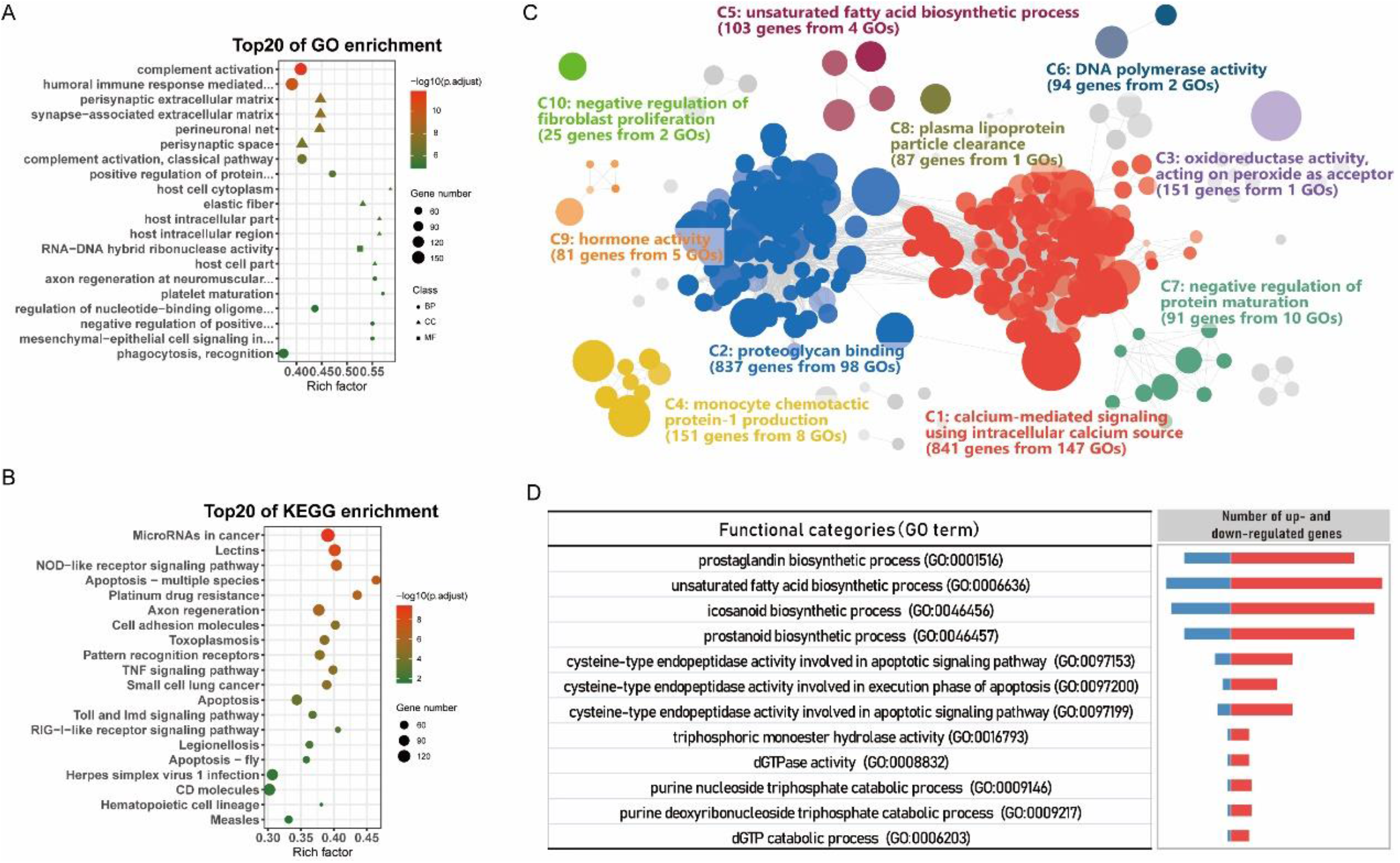
The top 20 significantly enriched GO terms for DEGs. **Fig.2B** The top 20 significantly enriched KEGG pathways for DEGs. **Fig.2C** Functional clusters enriched among DEGs. **Fig.2D** Number of different expression genes in clusters5, 19, and 25.

To analyze the differences between the two lines more comprehensively, we identified 26 functional clusters using GOMCL (Wang et al. 2020) (g-1). In short, GOMCL clustered enriched GO terms that had shared genes (>95%) using Markov Clustering and identified non-redundant representative functions within a GO network. The top ten root clusters included over 90.1% of the GO-annotated DEGs (Figure 2C). This approach revealed that calcium-mediated signaling using intracellular calcium sources (C1) and proteoglycan binding (C2), which are closely related to the neuroendocrine-immune system, accounts for the largest proportion (76.6%) among DEGs between two lines of oysters. Moreover, other functional clusters are associated with lipid metabolism and transport (C5, 8, and 24), DNA replication and repair (C6, 12, and 23), cell proliferation (C10) and apoptotic (C16 and 19), and tetrapyrrole catabolic process (C18), which shows the complex and diverse differences between two oyster strains.

What’s more, we examined all genes represented in 26 clusters. Interestingly, there were many more up-regulated genes than repressed genes in most clusters with significant differences (Table S5-2), especially in unsaturated fatty acid biosynthetic process (C5), cysteine-type endopeptidase activity involved in apoptotic process (C19), and purine nucleoside triphosphate catabolic process (C25). Further, we showed the gene expression profile of GO terms in these three clusters (Figure 2D), and all GO terms had more genes induced than repressed when comparing HH lines with CC, which may imply the intensely perturbing of energy metabolism, lipid metabolism, and apoptosis process in the orange-shell line. And, there were 21 DEGs were associated with the above GO terms and ranked in the top 10% of the significance of DEGs (Table 1).

**Table 1.** Significantly differentially expressed protein-coding transcripts.

| Transcript ID | log2FoldChange | padj | gene name |
| --- | --- | --- | --- |
| XM_034455638.1 | 3.791493806 | 6.43E-72 | caspase-7 |
| XM_011442211.3 | 6.160331262 | 2.02E-55 | cytochrome P450 1A1-like |
| XM_034459715.1 | -2.782676459 | 7.54E-50 | adenosine deaminase |
| XM_034479158.1 | 17.39844081 | 1.02E-42 | allene oxide synthase-lipoxygenase protein |
| XM_034455744.1 | -5.454240504 | 1.29E-39 | annexin A7-like |
| XM_034448556.1 | 6.690330785 | 1.07E-38 | caspase-14-like |
| XM_034459721.1 | 16.0020048 | 4.08E-38 | adenosine deaminase |
| XM_034476156.1 | 14.12484421 | 1.70E-28 | trichohyalin |
| XM_034462806.1 | 13.97698522 | 3.40E-28 | deoxynucleoside triphosphate triphosphohydrolase SAMHD1 |
| XM_034461487.1 | 2.115285699 | 8.01E-28 | 15-hydroxyprostaglandin dehydrogenase [NAD(+)] |
| XM_034465032.1 | 14.09964555 | 5.65E-27 | deoxynucleoside triphosphate triphosphohydrolase SAMHD1-like |
| XM_034465031.1 | 13.66416216 | 8.65E-27 | deoxynucleoside triphosphate triphosphohydrolase SAMHD1-like |
| XM_034479160.1 | 13.38994169 | 1.58E-25 | arachidonate 8S-lipoxygenase-like |
| XM_011441469.3 | 1.718474624 | 4.64E-25 | vitamin D3 receptor |
| XM_034454838.1 | -1.839185712 | 1.08E-22 | eosinophil peroxidase-like |
| XM_034450363.1 | 13.32825367 | 1.27E-22 | uncharacterized LOC105317447 |
| XM_034476154.1 | 13.14640311 | 3.88E-22 | trichohyalin |
| XM_011455884.3 | 2.115970827 | 1.17E-21 | steroid 17-alpha-hydroxylase/17,20 lyase |
| XM_034479841.1 | 14.07365518 | 2.81E-20 | deoxynucleoside triphosphate triphosphohydrolase SAMHD1-like |
| XM_034448595.1 | -2.35097917 | 5.39E-20 | steroid 17-alpha-hydroxylase/17,20 lyase-like |
| XM_034476157.1 | 12.30428867 | 1.21E-19 | trichohyalin |

### Analysis of differentially expressed lncRNA

We identified 2677 differential expression lncRNAs in the HH lines compared to CC, including 1467 that were down-regulated 54.8%) and 1210 that were up-regulated (45.2%) lncRNAs (Fig S2B and Table S2-2). The prediction of cis-and trans-target DEGs for DELs was used to understand the function of the lncRNAs. In short, with co-expression and co-localization analyses of strict filtering criteria, we identified 1784 cis-target mRNAs and 1983 trans-target DEGs from DELs (Table S6).

Then, all target genes were subjected to GO and KEGG pathway enrichment analysis. GO analysis of the target genes regulated by DELs revealed 288 significantly enriched terms (28 under molecular function, 18 under cellular component, and 242 under biological process, Table S7-1). As mentioned above, we identified 12 functional clusters by using GOMCL, of which the top ten largest clusters represented 98.4% of the total number of genes from all clusters (Table S7-2 and Fig.3A). The largest GO terms proportion among all clusters was related to immune response and protein K63-linked ubiquitination. According to some reports, K63-ubiquitination plays an important role in NF-κB activation, inflammation, and AKT kinase activation (Guocan et al. 2012). Correspondingly, the target genes were enriched in 20 pathways (Table S7-2), and these signaling pathways were related to apoptosis and some immune-related pathways, such as lectins, NOD-like receptor signaling pathway, RIG-I-like receptor signaling pathway, and NF-kappa B signaling pathway.

**Fig.3 A.**
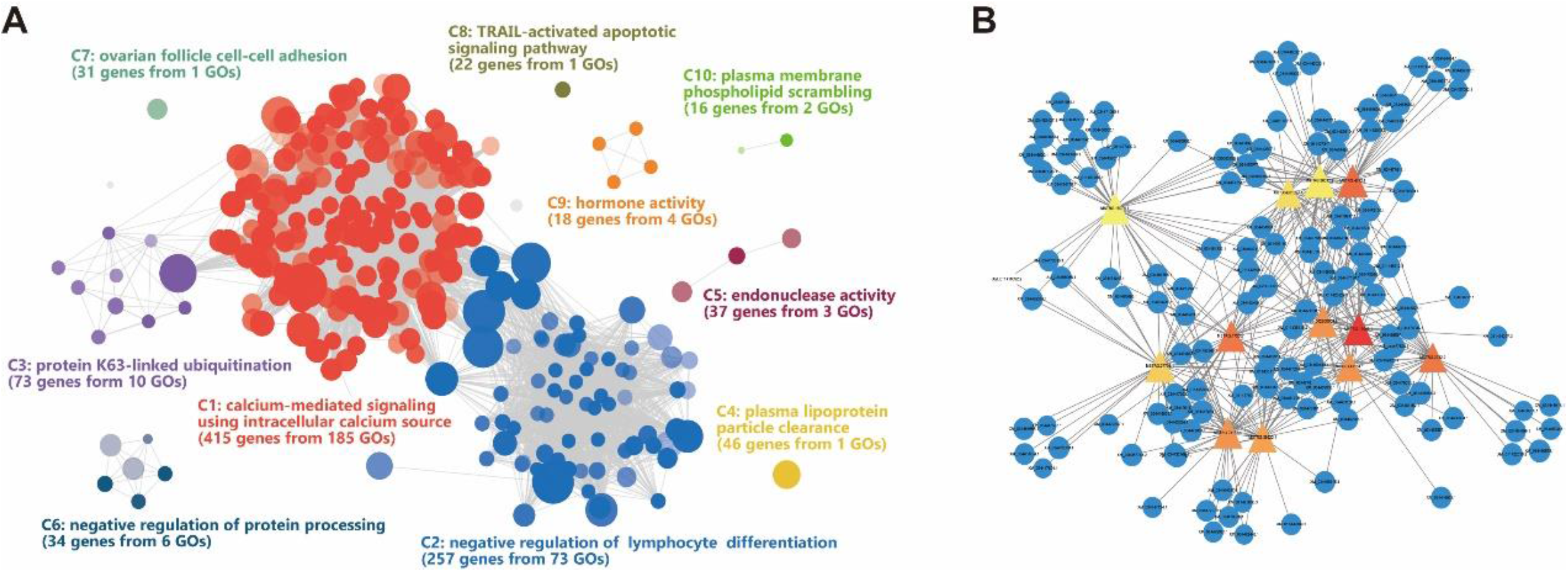
Functional clusters enriched among DELs. **Fig. 3B** The DELs-DEGs network of hub-DELs (degree > 30).

Hub has been considered that play essential roles in biological networks as they have high connectivity and are critical for maintaining the stability of the network. Thus, we analyzed the hub-DELs, which rank at the highest degree of nodes (degree > 30) in the DELs-DEGs network (Fig.3B). The result shows that these hub-DELs have extremely significant differences in expression between the two lines, and we detected some target genes function involvement of cell apoptosis, immune response, and lipid homeostasis (Table 2).

**Table 2.** hub-DELs and part of its target gene.

| Hub-DELs ID | log2FC | FDR | Degree | part of target gene |
| --- | --- | --- | --- | --- |
| MSTRG.15008.1 | 8.64 | 9.80E-52 | 45 | Cd209e(LOC117687969), GIMAP5(LOC117690697), RBM25(LOC105332454), XKR6(LOC117686702) |
| MSTRG.4742.1 | 12.21 | 3.77E-22 | 41 | C1QL4(LOC105327058), Cd209e(LOC117687969), ced-1(LOC117684820), NEURL1B(LOC105343065), RBM25(LOC105332454) |
| MSTRG.2732.2 | 5.04 | 4.83E-22 | 40 | ANGPT4(LOC105320234), RBM25(LOC105332454) |
| MSTRG.17565.1 | 2.97 | 1.33E-20 | 40 | GIMAP5(LOC117690697), TRIM56(LOC105330836) |
| MSTRG.41164.1 | 3.61 | 1.34E-20 | 38 | RBM25(LOC105332454), Notch2(LOC105348149) |
| MSTRG.24155.1 | 8.69 | 3.57E-16 | 38 | ANGPT4(LOC105320234), C1QB(LOC117690058), Colec11(LOC105322478), MALRD1(LOC117684818), Notch2(LOC105348149), TRIM56(LOC105330836) |
| XR_002200804.2 | 2.75 | 1.06E-13 | 38 | C1QB(LOC117690058), Cd209e(LOC117687969), Colec11(LOC105322478), CUBN(LOC105333487), GIMAP5(LOC117690697), MALRD1(LOC117684818), TRIM56(LOC105330836) |
| MSTRG.39523.1 | 6.40 | 3.68E-12 | 37 | C1QB(LOC117690058), Colec11(LOC105322478), MALRD1(LOC117684818), Notch2(LOC105348149), TRIM56(LOC105330836) |
| MSTRG.27708.1 | 4.07 | 3.13E-11 | 34 | ANGPT4(LOC105320234), Colec11(LOC105322478), MALRD1(LOC117684818), TRIM56(LOC105330836) |
| XR_004601027.1 | 4.01 | 1.52E-08 | 33 | C1QL4(LOC105327058), Cd209e(LOC117687969), GIMAP5(LOC117690697), NEURL1B(LOC105343065) |
| MSTRG.26017.9 | 8.30 | 1.62E-06 | 32 | C1QL4(LOC105327058), Cd209e(LOC117687969), ced-1(LOC117684820), GIMAP5(LOC117690697), NEURL1B(LOC105343065), RBM25(LOC105332454) |
| MSTRG.16974.1 | 8.07 | 1.02E-05 | 31 | ANGPT4(LOC105320234), P2RX7(LOC117688464) |

### Analysis of differentially expressed microRNA

Here, we identified 28 differential expression microRNAs (16 upregulated and 12 downregulated miRNAs, Fig S2C and Table S2-3) in the HH lines compared to CC. A total of 487 mRNAs were predicted as potential targets (table S8). We performed pathway enrichment analysis of these target genes to identify the biological functions of DEMs. The KEGG results revealed the enriched pathways involved in nuclear receptors, NK cell-mediated cytotoxicity, TNF signaling, p53 signaling, apoptosis, MAPK signaling, serotonergic synapse, and AGE-RAGE signaling in diabetic patients complications (table S9).

To increase our understanding of the molecular regulation mechanism of oyster lines difference, based on the data for DEGs, DEMs, and DELs, we used three software RNAhybrid (v2.1.2), Miranda (v3.3a), and TargetScan (v7.0) to identify biological targets of each DEMs from the DEGs and DELs that showed a significantly negative correlation with DEMs expression, subsequently obtained the DEGs-DEMs and DELs-DEMs pairs, then constructed the competing endogenous RNA (ceRNA) network (Fig.4A). As a result, 90 DELs, 22 DEMs, and 248 DEGs with interactive relationships were identified. Furth, we used the Spearman rank correlation coefficient (SCC) to assess correlations of expression between the DEGs-DEMs and DELs-DEMs pairs. Pairs with SCC < −0.6 and p-value < 0.01 were selected. Meanwhile, DELs-DEGs pairs with PCC > 0.9 and p-value < 0.01 were selected. Finally, we generated a high confidence ceRNA networks, which may play a key role in regulating molecular mechanisms of inbreeding depression (Fig.4B).

**Fig.4 A.**
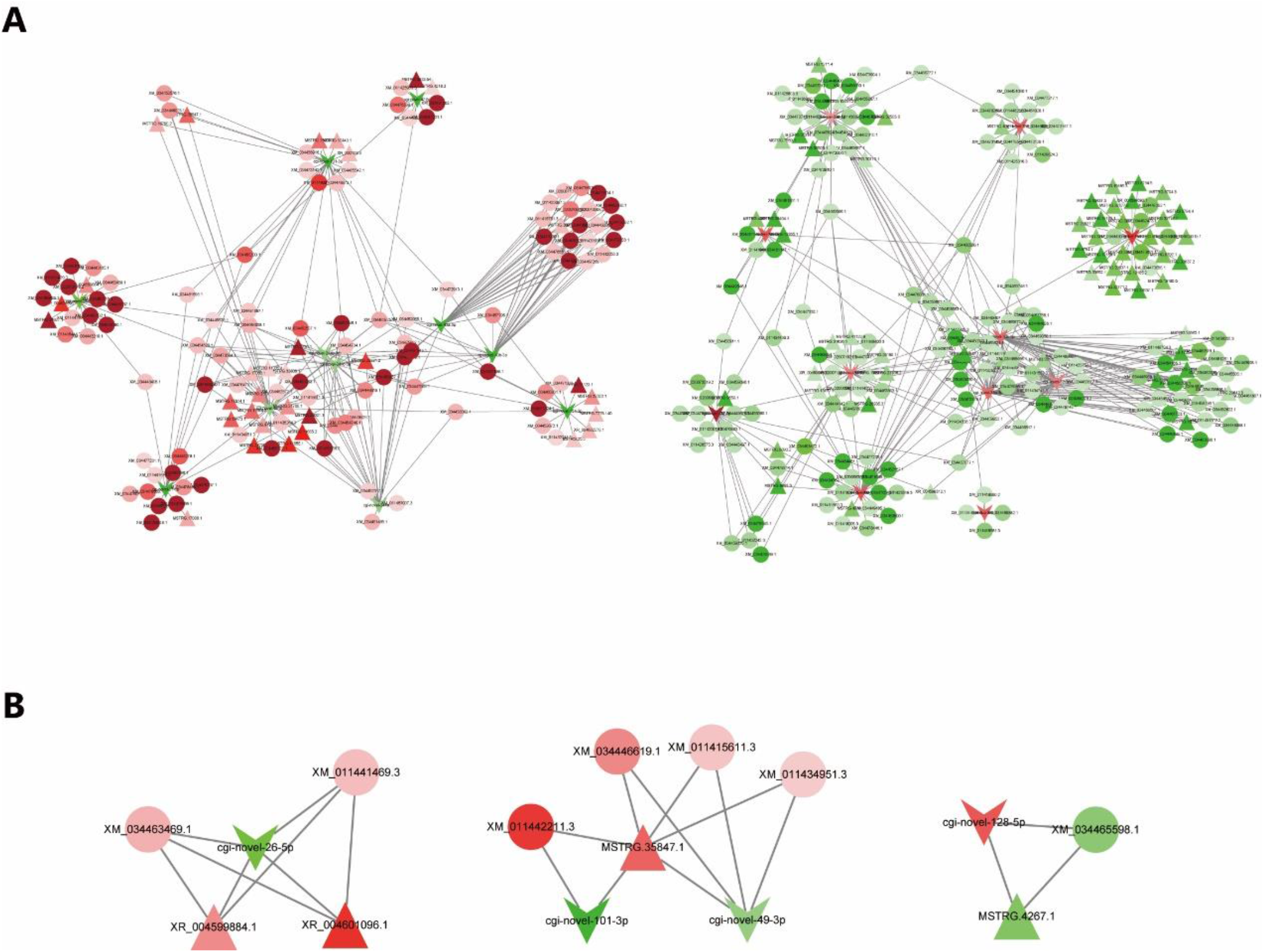
An overview of the competing endogenous RNA (ceRNA) network. Rectangle, ellipse and V indicate miRNA, protein-coding transcript and lncRNA transcript, respectively. Green and red indicate down-and up-regulation, respectively. **Fig. 4B** Predicted interaction of a high confidence ceRNA networks in regulating of inbreeding depression.

## Discussion

In this study, we used strand-specific RNA-seq and small RNA-seq data from two lines of the pacific oyster to provide a systematic view of the whole transcriptome level changes that occur within the inbreeding depression. By applying a set of stringent lncRNA and miRNAs identify pipelines, we identified 6,350 putative lncRNAs and 376 miRNAs. Moreover, the characteristics of lncRNAs and miRNAs were consistent with previous reports in oysters, which suggests that the ncRNAs identified were reliable. We believe this identified data about lncRNAs and miRNAs will serve as a better reference for related analysis in the oysters.

The characterized of inbreeding depression is often regarded as reduced individual fertility of offspring and a general reduction in fitness over environments. Although the study related to the molecular mechanism of inbreeding depression is necessarily limited to a small number of traits, territories, and populations, several studies of inbreeding and inbreeding depression found a tremendous amount of genes significant response with inbred, which function associated with immunity, stress, and metabolism(Kristensen et al. 2005; Ayroles et al. 2009; Zhao et al. 2019). Similarly, with the analysis of the expression patterns of mRNAs, we revealed enormous and multiple differences in genes transcript, and the result showed that the largest proportion of DEGs is related to the neuroendocrine-immune system. Meanwhile, we found a fascinating phenomenon that the majority of transcripts which function related to unsaturated fatty acid biosynthetic process, cysteine-type endopeptidase activity involved in the apoptotic process, and purine nucleoside triphosphate catabolic process were down-regulated in the orange-shell line. Some of these responding gene known to be involved in regulate immune system, for example SAMHD1 (LOC105317615), SAMHD1-like (LOC117692317), ADA (LOC105325902), CYP1A1-like (LOC105337485), allene oxide synthase-lipoxygenase protein gene (LOC105333406), HPGD (LOC105336455), and Alox8-like (LOC117692116) are involved in the pathway of eicosanoid synthesis and/or degradation, which relate to inflammation, stimulation of lipid metabolism, and regulation of insulin sensitivity (Nebert and Karp 2008). What’s more, VDR (LOC105336956) may affect shell biomineralization by mediating the action of vitamin D3 on cells (Zheng et al. 2019). The results indicated that these genes abnormally expressed may be one of the reasons for inbreeding depression of the orange line.

Epigenetics to the differential regulation of gene expression and silencing may shed fresh light on the impact of inbreeding depression (Biémont 2010). Researches about the function of ncRNAs in inbreeding depression were barely reported previously, especially in aquaculture species. Our results provide novel information regarding the potential regulatory roles of lncRNAs and miRNAs in inbred.

A mass of DELs was identified between two oyster lines, indicating that lncRNAs may have an essential regulating function in inbreeding depression. And enrichment analysis shows that most DELs regulate the gene related to the immune response process. We further analyzed the hub-DELs in the DELs-DEGs network. The result indicates that DELs directly regulate some key genes, which affect cell apoptosis, immune response, and lipid homeostasis. Although the role of these lncRNAs in regulating function requires further experimental validation, this information may help us explore the underlying mechanisms by which lncRNAs regulate inbreed depression in rare breeds.

Accumulating evidence indicates that miRNAs are an essential part of the complex regulatory networks that control cellular processes (Momen-Heravi and Bala 2018). Previous studies have taken advantage of the phenotypic and genetic variations across inbred strains of model animals to understand the regulation function of miRNAs connected with complex behavior and disease (Parsons et al. 2008). we identified 28 differential expression microRNAs (16 upregulated and 12 downregulated miRNAs) in the HH lines compared to CC. Although some miRNAs have high conservation between species (Moran et al. 2017), the DEMs identify in this study are all novel, which may be the unique miRNA in oysters or have an unknown function in inbreeding depression. Thus, we revealed the potential function of a few DEMs in inbreeding depression through pathway analysis of target genes. Many of DEMs are related to some signaling pathways which involved in regulating the cell proliferation, differentiation, apoptosis, and modulation of immune responses and induction of inflammation. Our results provide novel information regarding the regulatory roles of these miRNAs.

The competing endogenous RNA (ceRNA) hypothesis postulates that an increase in the cellular concentration of miRNA target RNA would reduce the cytoplasmic availability of the specific miRNA by binding it, thereby derepressing other mRNAs that are targets of the same miRNA (Gebert and MacRae 2019). Through interaction analysis of DEGs, DELs, and DEMs, we constructed a ceRNA relationship network by integrating multiple omics analyses. Furth, correlation analysis for the RNA pairs in the constructed ceRNA network revealed that several networks are conformed to the ceRNA hypothesis and detected ncRNA might be relevant to inbreeding depression in oysters. The genes associated with neuronal viability and polarity (Y. Chen et al. 2020; Jensen et al. 2000), such as WDR47 and NOVA1, could be modulated by cgi-novel-49-3p and MSTRG.35847.1. Megf10 is an epidermal growth factor (EGF) repeat-containing transmembrane protein involved in phagocytosis, astrocytes (Iram et al. 2016), and plays a role in muscle cell proliferation (Saha et al. 2017), has a relatively lower expression in the HH of CC and was found to be under the control of the MSTRG.4267.1 and cgi-novel-128-5p. What’s more, CYP1A1-like and VDR genes are also constructed by ceRNA network, which may have a key regulation function in inbreeding depression, as mentioned above. These provide direct evidence that ncRNAs, as regulators of gene expression, could modulate multiple subsystems or potential essential genes involved in inbreeding depression.

## Conclusion

In summary, we conducted a whole-transcriptome study on two strains of the pacific oyster with similar genetic backgrounds but different genetic variations and identified lncRNAs and miRNAs by stringent bioinformatics pipelines. And we identified and characterized DEGs, DELs, and DEMs that were involved in inbreeding depression. We also constructed potential regulatory networks of the molecular mechanisms of inbreeding depression. Our study enriches the expression profile of ncRNAs in the oyster. It provides valuable resources for further understanding the genetic basis of regulation of bivalve inbreeding depression from epigenetics. Such information potentially contributes to the understanding regulation mechanism of inbreeding depression and benefits the genetic breeding in rare breeds.

## Author Contributions

SL conceived the study and obtained the funding. ZF performed the experiment. SC analyzed the related data, drafted the manuscript and produced figures and tables. SL revised the manuscript. QL supervised the work. All authors have read and approved the final manuscript.

## Data Availability Statement

The datasets for this study can be found in the NCBI Sequence Read Archive (SRA) accession:

## Reference

Ayroles JF, Hughes KA, Rowe KC, Reedy MM, Rodriguez-Zas SL, Drnevich JM, Cáceres CE, Paige KN (2009) A Genomewide Assessment of Inbreeding Depression: Gene Number, Function, and Mode of Action. Conservation Biology 23:920–930. https://doi.org/10.1111/j.1523-1739.2009.01186.x.

Biémont C (2010) Inbreeding Effects in the Epigenetic Era. Nature Reviews Genetics 11:234. https://doi.org/10.1038/nrg2664-c1.

Bolger AM, Lohse M, Usadel B (2014) Trimmomatic: A Flexible Trimmer for Illumina Sequence Data. Bioinformatics 30:2114–2020. https://doi.org/10.1093/bioinformatics/btu170.

Bowley, SC, Comizzoli P, Lindell KA, Matsas DJ, White EC (2019) Genetic Cryopreservation of Rare Breeds of Domesticated North American Livestock: Smithsonian & SVF Biodiversity Preservation Project. Diversity 11:198. https://doi.org/10.3390/d11100198.

Buchfink B, Xie C, Huson DH (2015) Fast and Sensitive Protein Alignment Using DIAMOND. Nature Methods 12:59–60. https://doi.org/10.1038/nmeth.3176.

Chen S, Zhou Y, Chen Y, Gu J (2018) Fastp: An Ultra-Fast All-in-One FASTQ Preprocessor. Bioinformatics 34(17):i884–i890. https://doi.org/10.1093/bioinformatics/bty560.

Danchin-Burge C, Palhière I, François D, Bibé B, Leroy G, Verrier E (2010) Pedigree Analysis of Seven Small French Sheep Populations and Implications for the Management of Rare Breeds. Journal of Animal Science 88:505–516. https://doi.org/10.2527/jas.2009-1961.

Enright AJ, John B, Gaul U, Tuschl T, Sander C, Marks DS (2003) MicroRNA Targets in Drosophila. Genome Biology 4(11): 1–27. http://genomebiology.com/2003/5/1/R1

Fang J, Han Z, Li Q (2021) Effect of Inbreeding on Performance and Genetic Parameters of Growth and Survival Traits in the Pacific Oyster Crassostrea Gigas at Larval Stage. Aquaculture Reports 19: 100590. https://doi.org/10.1016/j.aqrep.2021.100590.

Friedländer MR, Mackowiak SD, Li N, Chen W, Rajewsky N (2012) MiRDeep2 Accurately Identifies Known and Hundreds of Novel MicroRNA Genes in Seven Animal Clades. Nucleic Acids Research 40:37–52. https://doi.org/10.1093/nar/gkr688.

Han Z, Li Q (2018) Different Responses between Orange Variant and Cultured Population of the Pacific Oyster Crassostrea Gigas at Early Life Stage to Temperature-Salinity Combinations. Aquaculture Research 49:2233–2239. https://doi.org/10.1111/are.13680.

Han Z, Li Q, Liu S, Yu H, Kong L (2019) Genetic Variability of an Orange-Shell Line of the Pacific Oyster Crassostrea Gigas during Artificial Selection Inferred from Microsatellites and Mitochondrial COI Sequences. Aquaculture 508:159–166. https://doi.org/10.1016/j.aquaculture.2019.04.074.

Kaikkonen MU, Lam MTY, Glass CK (2011) Non-Coding RNAs as Regulators of Gene Expression and Epigenetics. Cardiovascular Research 90:430–440. https://doi.org/10.1093/cvr/cvr097.

Kang YJ, Yang DC, Kong L, Hou M, Meng YQ, Wei L, Gao G (2017) CPC2: A Fast and Accurate Coding Potential Calculator Based on Sequence Intrinsic Features. Nucleic Acids Research 45:12–16. https://doi.org/10.1093/nar/gkx428.

Kim D, Paggi JM, Park C, Bennett C, Salzberg SL (2019) Graph-Based Genome Alignment and Genotyping with HISAT2 and HISAT-Genotype. Nature Biotechnology 37:907–915. https://doi.org/10.1038/s41587-019-0201-4.

Kristensen TN, Sørensen P, Kruhøffer M, Pedersen KS, Loeschcke V (2005) Genome-Wide Analysis on Inbreeding Effects on Gene Expression in Drosophila Melanogaster. Genetics 171:157–167. https://doi.org/10.1534/genetics.104.039610.

Kruger J, Rehmsmeier M (2006) RNAhybrid: MicroRNA Target Prediction Easy, Fast and Flexible. Nucleic Acids Research 34:451–454. https://doi.org/10.1093/nar/gkl243.

Langmead B, Trapnell C, Pop M, Salzberg SL (2009) Ultrafast and Memory-Efficient Alignment of Short DNA Sequences to the Human Genome. Genome Biology 10:R25. https://doi.org/10.1186/gb-2009-10-3-r25.

Lewis BP, Burge CB, Bartel DP (2005) Conserved Seed Pairing, Often Flanked by Adenosines, Indicates That Thousands of Human Genes Are MicroRNA Targets. Cell 120:15–20. https://doi.org/10.1016/j.cell.2004.12.035.

Li A, Zhang J, Zhou Z (2014) PLEK: A Tool for Predicting Long Non-Coding RNAs and Messenger RNAs Based on an Improved k-Mer Scheme. BMC Bioinformatics 15:311. https://doi.org/10.1186/1471-2105-15-311.

Love MI, Huber W, Anders S (2014) Moderated Estimation of Fold Change and Dispersion for RNA-Seq Data with DESeq2. Genome Biology 15:550. https://doi.org/10.1186/s13059-014-0550-8.

Marc C, Herve P (2017) AnnotationForge. Bioconductor. https://doi.org/10.18129/B9.BIOC.ANNOTATIONFORGE.

Martin M (2011) Cutadapt Removes Adapter Sequences from High-Throughput Sequencing Reads. EMBnet.Journal 17:10. https://doi.org/10.14806/ej.17.1.200.

Mistry J, Finn RD, Eddy SR, Bateman A, Punta M (2013) Challenges in Homology Search: HMMER3 and Convergent Evolution of Coiled-Coil Regions. Nucleic Acids Research 41:e121. https://doi.org/10.1093/nar/gkt263.

Molnár J, Nagy T, Stéger V, Tóth G, Marincs F, Barta E (2014) Genome Sequencing and Analysis of Mangalica, a Fatty Local Pig of Hungary. BMC Genomics 15:761. https://doi.org/10.1186/1471-2164-15-761.

Nandolo W, Mészáros G, Banda LJ, Gondwe TN, Lamuno D, Mulindwa HA, Nakimbugwe HN, Wurzinger M, Utsunomiya YT, Woodward-Greene MJ, Liu M, Liu G, Van Tassell C P, Curik I, Rosen BD, Sölkner J (2019) Timing and Extent of Inbreeding in African Goats. Frontiers in Genetics 10:537. https://doi.org/10.3389/fgene.2019.00537.

Nebert DW, Karp CL (2008) Endogenous Functions of the Aryl Hydrocarbon Receptor (AHR): Intersection of Cytochrome P450 1 (CYP1)-Metabolized Eicosanoids and AHR Biology. Journal of Biological Chemistry 283:36061–36065. https://doi.org/10.1074/jbc.R800053200.

Paige KN (2010) The Functional Genomics of Inbreeding Depression: A New Approach to an Old Problem. BioScience 60:267–277. https://doi.org/10.1525/bio.2010.60.4.5.

Pertea G, Pertea M (2020) GFF Utilities: GffRead and GffCompare. F1000Research 9:304. https://doi.org/10.12688/f1000research.23297.2.

Pertea M, Pertea GM, Antonescu CM, Chang TC, Mendell JT, Salzberg SL (2015) StringTie Enables Improved Reconstruction of a Transcriptome from RNA-Seq Reads. Nature Biotechnology 33:290–295. https://doi.org/10.1038/nbt.3122.

Quinlan AR, Hall IM (2010) BEDTools: A Flexible Suite of Utilities for Comparing Genomic Features. Bioinformatics 26:841–842. https://doi.org/10.1093/bioinformatics/btq033.

The Aquaculture Genomics, Genetics and Breeding Workshop, Abdelrahman H, ElHady M, Alcivar-Warren A, Allen S, Al-Tobasei R, Bao L, et al (2017) Aquaculture Genomics, Genetics and Breeding in the United States: Current Status, Challenges, and Priorities for Future Research. BMC Genomics 18:191. https://doi.org/10.1186/s12864-017-3557-1.

Triantaphyllopoulos KA, Ikonomopoulos I, Bannister AJ (2016) Epigenetics and Inheritance of Phenotype Variation in Livestock. Epigenetics & Chromatin 9:31. https://doi.org/10.1186/s13072-016-0081-5.

Wang G, Yin H, Li B, Yu C, Wang F, Xu X, Cao J, Bao Y, Wang L, Abbasi AA, Bajic VB, Ma L, Zhang Z (2019) Characterization and Identification of Long Non-Coding RNAs Based on Feature Relationship. Bioinformatics 35:2949–2956. https://doi.org/10.1093/bioinformatics/btz008.

Wang G, Oh DH, Dassanayake M (2020) GOMCL: A Toolkit to Cluster, Evaluate, and Extract Non-Redundant Associations of Gene Ontology-Based Functions. BMC Bioinformatics 21:139. https://doi.org/10.1186/s12859-020-3447-4.

Wang G, Gao Y, Li R, Jin G, Cai Z, Chao JI, Lin HK (2012) K63-Linked Ubiquitination in Kinase Activation and Cancer. Frontiers in Oncology 2:5. https://doi.org/10.3389/fonc.2012.00005.

Wang L, Park HJ, Dasari S, Wang S, Kocher JP, Li W (2013) CPAT: Coding-Potential Assessment Tool Using an Alignment-Free Logistic Regression Model. Nucleic Acids Research 41:74. https://doi.org/10.1093/nar/gkt006.

Wu T, Hu E, Xu S, Chen M, Guo P, Dai Z, Feng T, Zhou L, Tang W, Zhan L, Fu X, Liu S, Bo X, Yu G (2021) ClusterProfiler 4.0: A Universal Enrichment Tool for Interpreting Omics Data. The Innovation 2:100141. https://doi.org/10.1016/j.xinn.2021.100141.

Xu F, Wang X, Feng Y, Huang W, Wang W, Li L, Fang X, Que H, Zhang G (2014) Identification of Conserved and Novel MicroRNAs in the Pacific Oyster Crassostrea Gigas by Deep Sequencing. PLoS ONE 9(8):e104371. https://doi.org/10.1371/journal.pone.0104371.

Zhao L, Li Y, Lou J, Yang Z, Liao H, Fu Q, Guo Z, Lian S, Hu X, Bao Z (2019) Transcriptomic Profiling Provides Insights into Inbreeding Depression in Yesso Scallop Patinopecten Yessoensis. Marine Biotechnology 21:623–633. https://doi.org/10.1007/s10126-019-09907-9.

Zheng Z, Hao R, Xiong X, Jiao Y, Deng Y, Du X (2019) Developmental Characteristics of Pearl Oyster Pinctada Fucata Martensii: Insight into Key Molecular Events Related to Shell Formation, Settlement and Metamorphosis. BMC Genomics 20:122. https://doi.org/10.1186/s12864-019-5505-8.

