## Supplementary figures and images for "Gene expression signatures underlying inbreeding depression as revealed by whole-transcriptome analysis of selectively bred strains of the Pacific Oyster (*Crassostrea gigas*)"

### Supplemental Figure S1

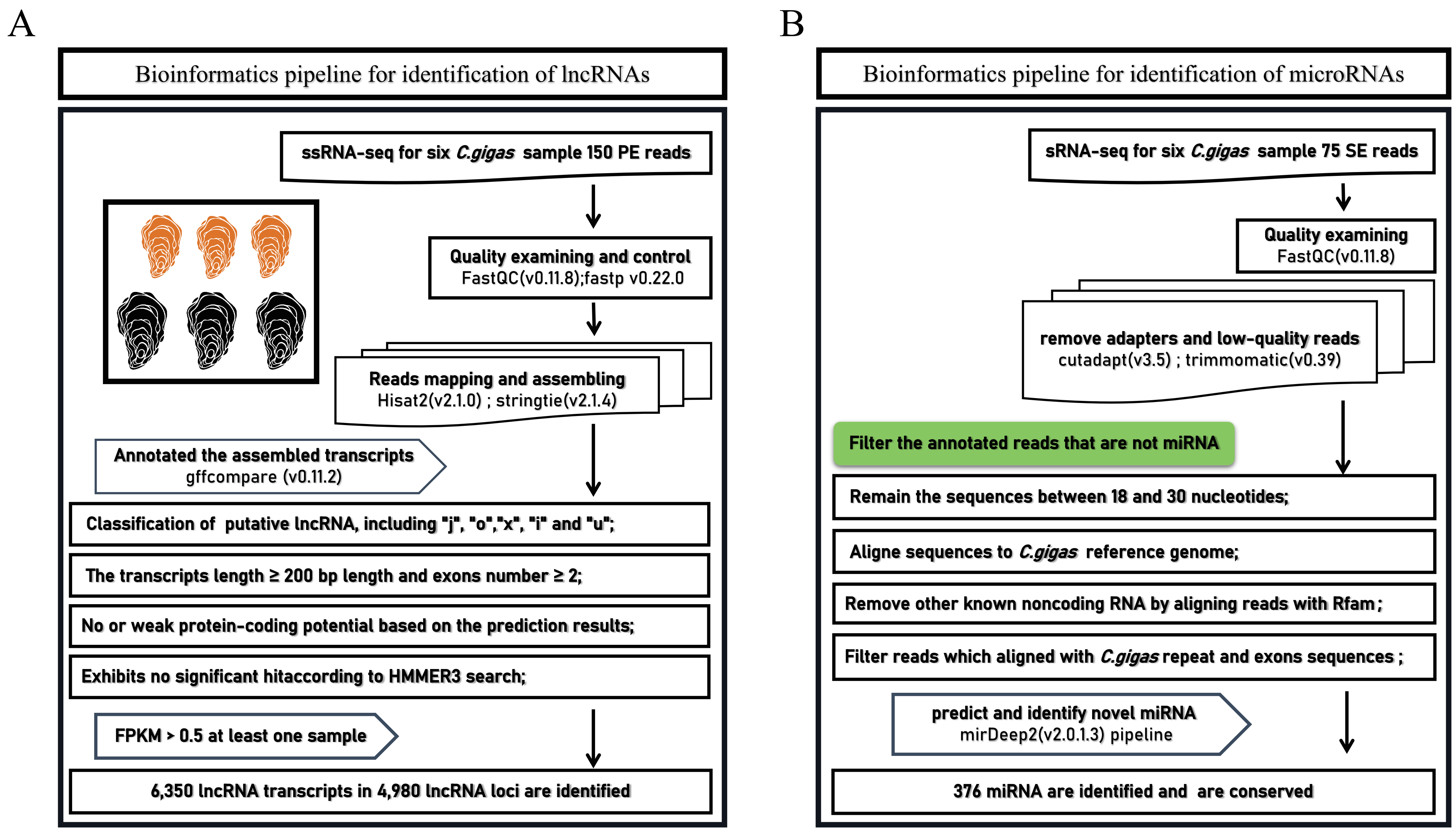

### Supplemental Figure S2A

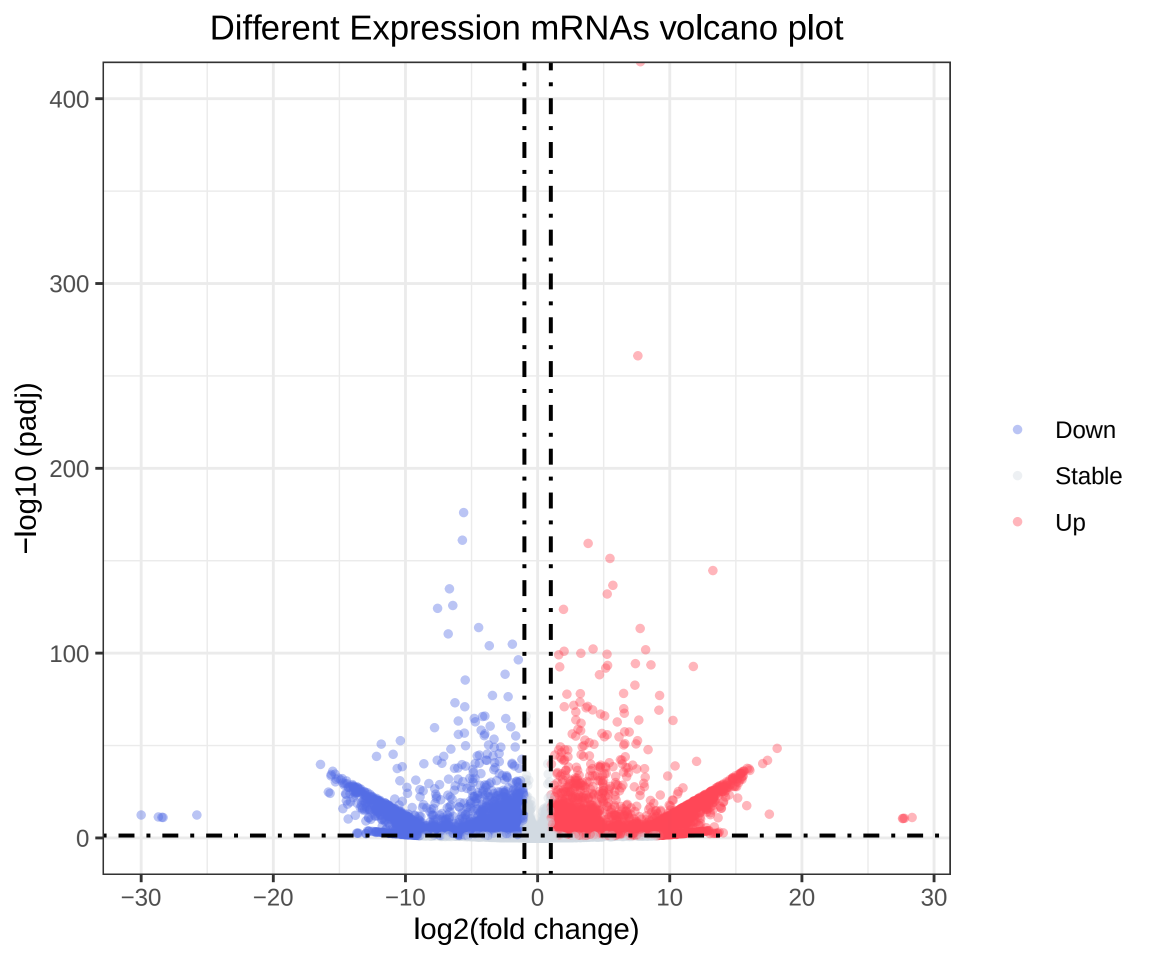

### Supplemental Figure S2B

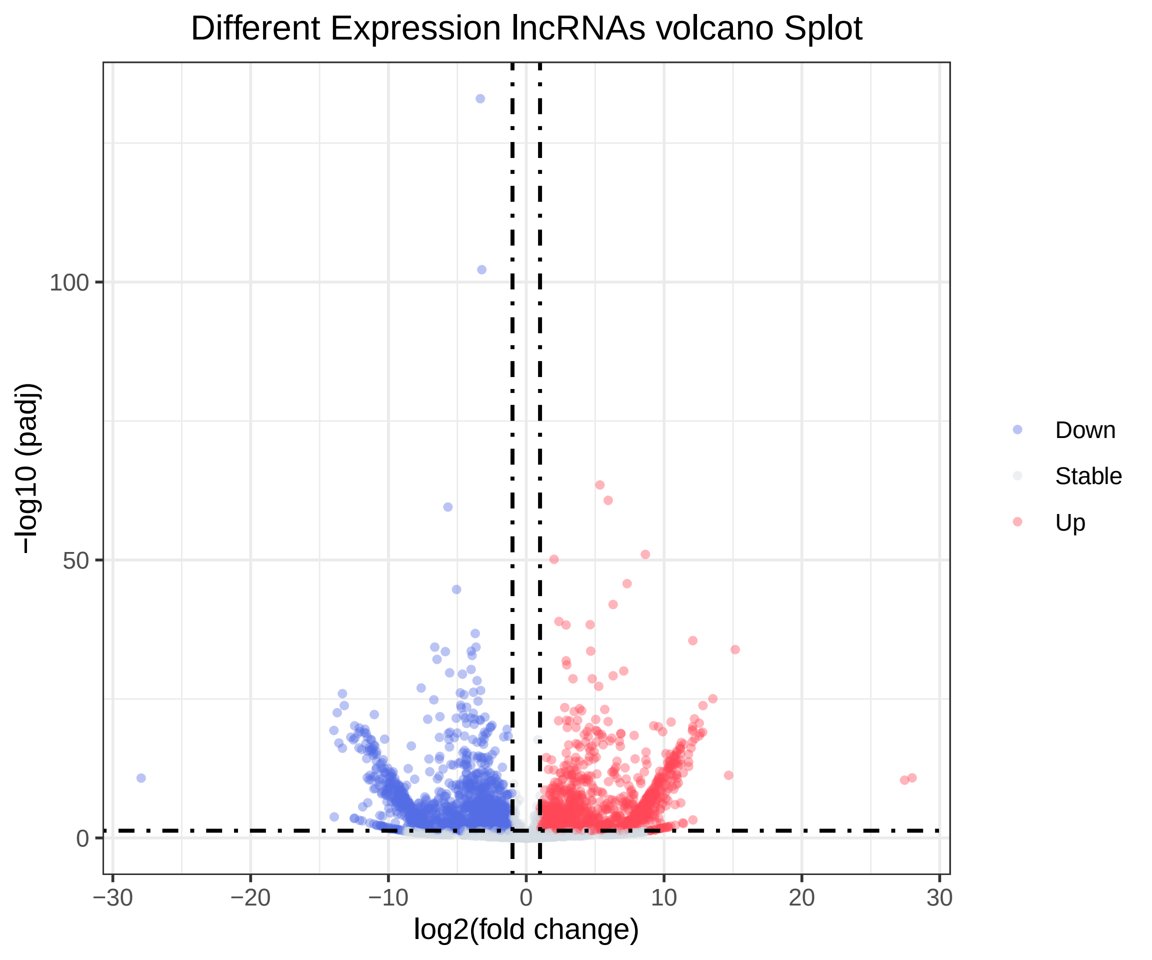

### Supplemental Figure S2C

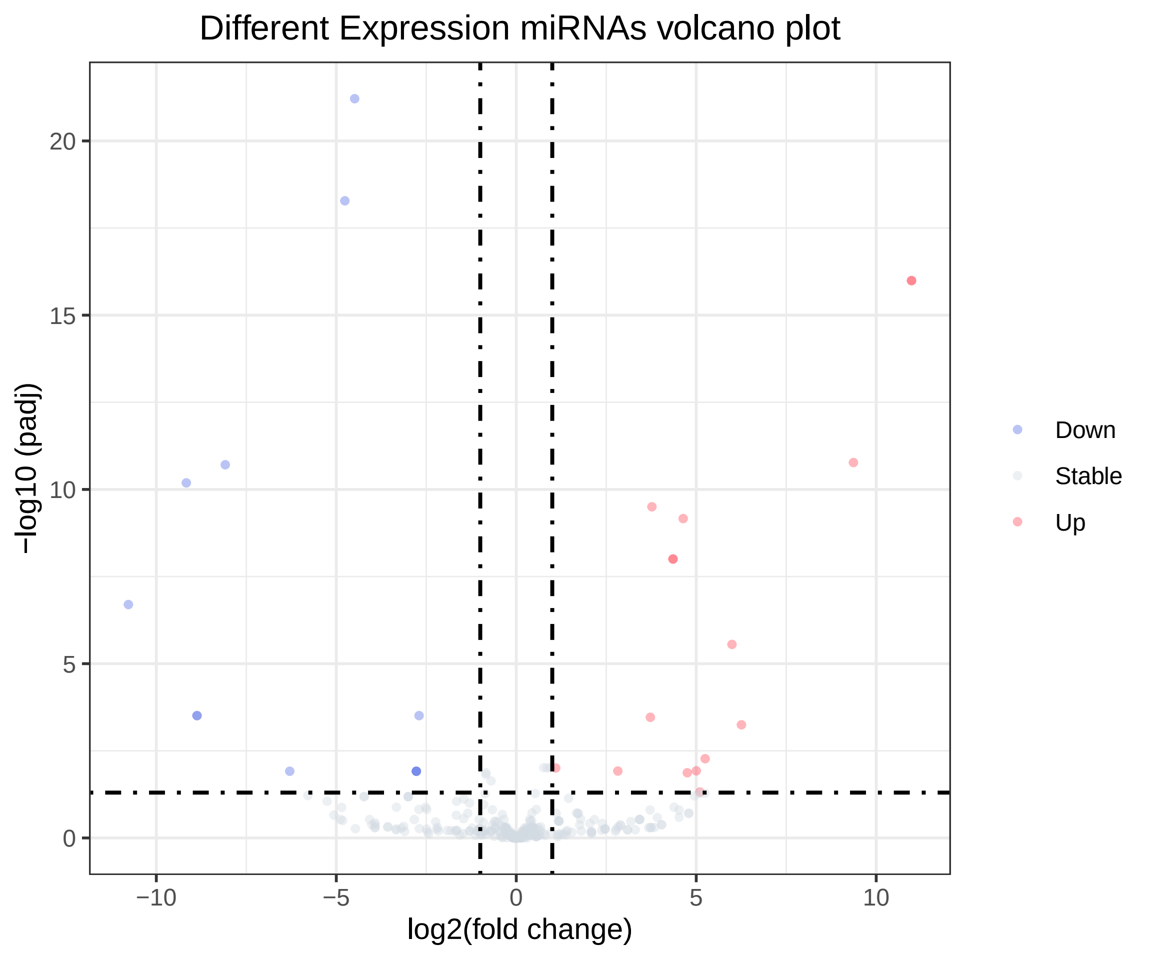
